# A method for rapid selection of randomly induced mutations in a gene of interest using CRISPR/Cas9 mediated activation of gene expression

**DOI:** 10.1101/788372

**Authors:** William A. Ng, Andrew Ma, Molly Chen, Bruce H. Reed

## Abstract

We have developed a CRISPR/Cas9 based method for isolating randomly induced recessive lethal mutations in a gene of interest (GOI) by selection within the F1 progeny of a single genetic cross. Our method takes advantage of the ability to overexpress a GOI using CRISPR/Cas9 mediated activation of gene expression. In essence, the screening strategy is based upon the idea that if overexpression of a wild type allele can generate a phenotype, then overexpression of a newly induced loss-of-function allele will lack this phenotype. As a proof-of-principle, we used this method to select EMS induced mutations of the Drosophila gene *hindsight* (*hnt*). From approximately 45,000 F1 progeny we recovered 8 new EMS induced loss-of-function *hnt* alleles that we characterized as an allelic series of hypomorphic mutations. This new method can, in theory, be used to recover randomly induced point mutants in a GOI and can be applied to any circumstance where CRISPR/Cas9 mediated activation of gene expression is associated with lethality or a visible phenotype.

## Introduction

CRISPR/Cas9 based genome editing represents a significant advance within the field of genetics. In general, co-expression of Cas9 and an engineered single guide RNA (sgRNA) forms a sequence homology-dependent endonuclease that creates DNA double-stranded breaks (DSBs) within a specified target sequence (Jinek *et al*. 2012), and this has also been successfully used in Drosophila (Bassett *et al*. 2013; Gratz *et al*. 2013). DSBs can be repaired by the cell’s endogenous DNA repair machinery, either through error-prone non-homologous end-joining (NHEJ), possibly resulting in mutations – small insertions or deletion, or through homology directed repair (HDR), which requires template DNA containing sequences homologous to the regions flanking the DSB. The CRISPR/Cas9 system is now routinely used to facilitate gene knockout and gene replacement strategies in various model genetic organisms (Ma and Liu 2015).

The CRISPR/Cas9 system is remarkable in that it allows the localization of a protein/RNA complex (Cas9 + sgRNA) to a specific sequence within the genome, as specified through a 17-20 nucleotide region of the sgRNA molecule. Being able to localize a protein to a specific DNA sequence opens up many new opportunities, and Cas9 has been modified to create several new tools and techniques. By mutating the catalytic sites of the Cas9 endonuclease, a catalytically inactive or “dead” Cas9 (dCas9) has been developed (Qi *et al*. 2013). Various dCas9 fusion proteins have subsequently been designed, and the dCas9 system can be used for a variety of new applications, including the activation or repression of gene transcription, site-specific chromatin modification, or visualization of specific chromosome sites in living cells - see (Pulecio *et al*. 2017; Adli 2018) for reviews.

For Drosophila researchers, prior to the development of CRISPR/Cas9, targeted gene knockouts or gene replacement by homologous recombination were both possible, but challenging and labor intensive endeavors – see (Bibikova *et al*. 2002; Liu *et al*. 2012) for reviews of pre-CRISPR/Cas9 techniques. CRISPR/Cas9 strategies for gene knockout and gene replacement are now common techniques for many Drosophila research groups. Building on the *GAL4*/*UAS* system of inducible gene expression, CRISPR/Cas9 approaches for tissue specific target gene knockdown or overexpression in Drosophila are now also possible; resources, including >1600 transgenic stocks expressing different sgRNAs made by the Drosophila RNAi Screening Center and Transgenic RNAi Project (DRSC/TRiP) are publically available – see (Bier *et al*. 2018) for overview of resources. In the case of targeted gene overexpression, the TRiP transgenic lines were designed to ubiquitously express two sgRNAs targeted to sequences upstream of a gene’s transcriptional start site (TSS) (Lin *et al*. 2015; Ewen-Campen *et al*. 2017). These lines are known as *TRiP-OE*, or *TOE* lines. Target gene overexpression is achieved by crossing a *TOE* line to a line that carries a *GAL4* driver of one’s choice and *UAS-dCas9*.*VPR*, which is dead-Cas9 fused to the VPR transcriptional activation domain. Other sgRNA lines known as *TRiP-KO*, or *TKO* lines, designed for gene knockout, ubiquitously express a single sgRNA targeted to the coding region of a GOI, and can be used to create germline or somatic mutations.

We are interested in the regulation and function of the gene *hindsight* (*hnt*), which is the Drosophila homolog of mammalian *Ras Responsive Element Binding Protein-1* (*RREB-1*) (Melani *et al*. 2008; Ming *et al*. 2013). Most mutations of *hnt* are recessive and embryonic lethal, and fail in the morphogenetic process of germ band retraction (Yip *et al*. 1997). In this study we confirm *hnt* overexpression using a *TRiP-OE sgRNA* with *GAL4* and *UAS-dCas9*.*VPR*. Moreover, by reducing the number of functional copies of *hnt*^*+*^ from two to one, we found that known loss-of-function *hnt* mutations heterozygous to a wild type allele (*hnt*/ *hnt*^*+*^) can suppress the *hnt* overexpression phenotype.

Subsequently, we developed a method for screening for new loss-of-function *hnt* alleles. Our method is applicable to any GOI provided two requirements can be satisfied: first, the GOI must be associated with an overexpression phenotype induced by *UAS- dCas9*.*VPR* expression under *GAL4* control along with expression of an appropriate sgRNA; and second, the overexpression phenotype must either be sensitive to the dosage of wild type alleles, or if such is not the case, a refractory allele (an allele that is unresponsive to dCas9.VPR mediated activation of gene expression, but otherwise wild type) must be available. We demonstrate that *TOE-sgRNA* expression with active Cas9 is an efficient way to recover refractory alleles.

We present the results of this new screening strategy in which randomly induced recessive lethal alleles are selected in an F1 visible screen. Following EMS treatment of a responsive line, and crossing to a refractory line that carries *TOE-sgRNA* + *GAL4* + *UAS-dCas9*.*VPR*, we screened for the absence of an overexpression phenotype and, in so doing, selected 8 new alleles of *hnt* from ∼45,000 F1 progeny over a period of 2-3 weeks.

The recovery of numerous mutant alleles in a GOI is potentially useful for analysis of mutation profiles, or for the rapid recovery of an allelic series for a GOI. The ability to overexpress mutant alleles using dCas9.VPR mediated activation of gene expression can also be useful in classifying alleles as hypomorphic or nullomorphic.

## Materials and Methods

### Drosophila stocks

All Drosophila cultures were raised on standard medium at 25°C under a 12 hour light/dark cycle regime, unless otherwise indicated. Most stocks used in this study were either obtained from the Bloomington *Drosophila* Resource Center (BDRC), or derived from stocks obtained from the BDRC. The eye specific *GAL4* driver was *GMR-GAL4* = *P*{*GAL4-ninaE*.*GMR*}*12* (BDRC #1104). The stock ubiquitously expressing sgRNA targeted to *hnt* upstream region was *TOE-GS00052* = *y sc* v sev*^*21*^; *P*{*y[+t7*.*7] v[+t1*.*8]*=*TOE*.*GS00052*}*attP40* (BDRC #67530). The dCas9 transcriptional activator used was *UAS-dCas9*.*VPR* = *y w\**; *Kr*^*If*^ */CyO*; *P*{*y[+t7*.*7] w[+mC]*=*UAS- 3xFLAG*.*dCas9*.*VPR*}*attP2* (BDSC #66562). RNAi knockdown of *hnt* was achieved using *UAS-hnt-RNAi* = *y sc* v sev*^*21*^; *P*{*y[+t7*.*7] v[+t1*.*8]=TRiP*.*HMS00894*}*attP2*/ *TM3, Sb* (BDRC #33943). The expression of active Cas9 in the germline used *GAL4-nos*.*NGT*; *UAS-Cas9*.*P2* = *w*\*; *P*{*w[+mC]=GAL4-nos*.*NGT*}*40*; *P*{*y[+t7*.*7]=UAS-Cas9*.*P2*}*attP2* (BDRC #67083). The temperature sensitive *hnt* hypomorphic allele used was *hnt*^*peb*^ = *peb v* (BDRC #80). The deletion of the *hnt* gene used was *Df(1)ED67*27 = *Df(1)ED6727, w*^*1118*^ *P*{*w[+mW*.*Scer\FRT*.*hs3]=3’*.*RS5+3*.*3’*}*ED6727*/*FM7h* (BDRC #8956).

The chromosome used in the mutagenesis screen was *w PBac{RB}e02388b* (Exelixis at Harvard medical School #e02388). Pre-existing *hnt* alleles used included *hnt*^*XE81*^, *hnt*^*1142*^, *hnt*^*308*^, and *hnt*^*345x*^ as previously described (Yip *et al*. 1997; Wilk *et al*. 2000; Reed *et al*. 2001; Farley *et al*. 2018). Confocal imaging of pupal eye made use of *Ubi-DE-cadherin-GFP* as previously described (Reed *et al*. 2004). Complementation crosses were facilitated by the transgenic line *peb*^*BAC-CH321-46J02*^ as described (Farley *et al*. 2018). *GAL4* > *UAS* based overexpression of *hnt* used *UAS-GFP-hnt* as previously described (Baechler *et al*. 2015). This line *UAS-GFP-Hnt*, however, resulted in pupal lethality in combination with *GMR-GAL4*. To achieve lower levels of *hnt* overexpression not resulting in pupal lethality, the *UAS-GFP-hnt* insertion was mobilized by crossing to the transposase line *Δ2-3* (BDRC #3629). The insertion line *UAS-GFP-hnt*^*J27*^ was recovered and found to be viable in combination with the same *GMR-GAL4* driver.

### EMS Mutagenesis

3-5 day old males of the *w PBac{RB}e02388b* stock were collected and starved for 2 hours by placing in empty vials. Following starvation, male flies were transferred to bottles containing Kimwipe tissues soaked in a 2% sucrose solution containing 25 mM EMS (Sigma-Aldrich) and allowed to feed overnight. They were subsequently placed in vials with 3-5 day old virgin *RGV* females in crowded conditions and allowed to mate for 6-8 hours (30-40 virgin females with same number of males). Mated females and males were transferred to bottles for two days at which point they were removed.

### Molecular Characterization of Refractory Mutants

In order to characterize refractory alleles we extracted genomic DNA from a pooled group of 20-30 isogenic male flies using gSYNC DNA Extraction Kits (Geneaid) according to supplier’s instructions. We used 2.5 μL of isolated genomic DNA in 25 μL PCR reactions (2X FroggaMix; FroggaBio) utilizing primers flanking the guide target regions at ∼400 bp upstream and downstream of the sgRNA target sequences. PCR products were purified using GenepHlow Gel/PCR Kits (Geneaid) then sent out for Sanger sequencing using both forward and reverse primers.

### Complementation crosses

New *hnt* alleles, maintained as balanced lines over *FM7h*, were tested for complementation by crossing to *hnt*^*peb*^ males at the restrictive temperature 29°C as well as males of the following genotype: *y w hnt*^*XE81*^/*Y*; *peb*^*BAC-CH321-46J02*^/ *TM6C, Sb*. Failure to complement *hnt*^*peb*^ was indicated by a rough eye phenotype, while failure to complement *hnt*^*XE81*^ was indicated by the absence of *B*^*+*^ *Sb* female progeny carrying neither the *FM7h* balancer or the *peb*^*BAC*^ insertion.

### Immunostaining and Imaging

Immunostaining of embryos was carried out as described (Reed *et al*. 2001). Primary antibodies were used at the indicated dilutions: mouse monoclonal anti-Hindsight (Hnt) 27B8 1G9 (1:25; from H. Lipshitz, University of Toronto), guinea pig polyclonal anti-Hindsight (1:1000; from H. Lipshitz, University of Toronto). Secondary antibodies used were: Alexa Flour® 488 goat anti-mouse and TRITC goat anti-guinea pig (1:500; Cedarlane Labs). In most cases *hnt* mutant embryos were recognized by their u-shaped morphology, which is associated with a failure to complete germ band retraction and premature death of the amnioserosa.

Confocal microscopy and confocal image processing were performed as previously described (Cormier *et al*. 2012). Eye micrographs (Fig. 1A,B) were taken using a Nikon SMZ25 stereomicroscope equipped with Nikon Digital Sight Ri2 16.25MP colour camera and processed using extended-depth-of-focus feature of Nikon NIS-Elements Arv4.50 software. Other eye micrographs were acquired using a compound Olympus microscope with a 5X objective and oblique illumination. Z stacks were collected by manually adjusting the focal plane as images were collected using a mobile phone camera positioned over the ocular lens. Images collected were processed using ImageJ and an extended depth of focus plugin (Schneider *et al*. 2012).

**Figure 1.**
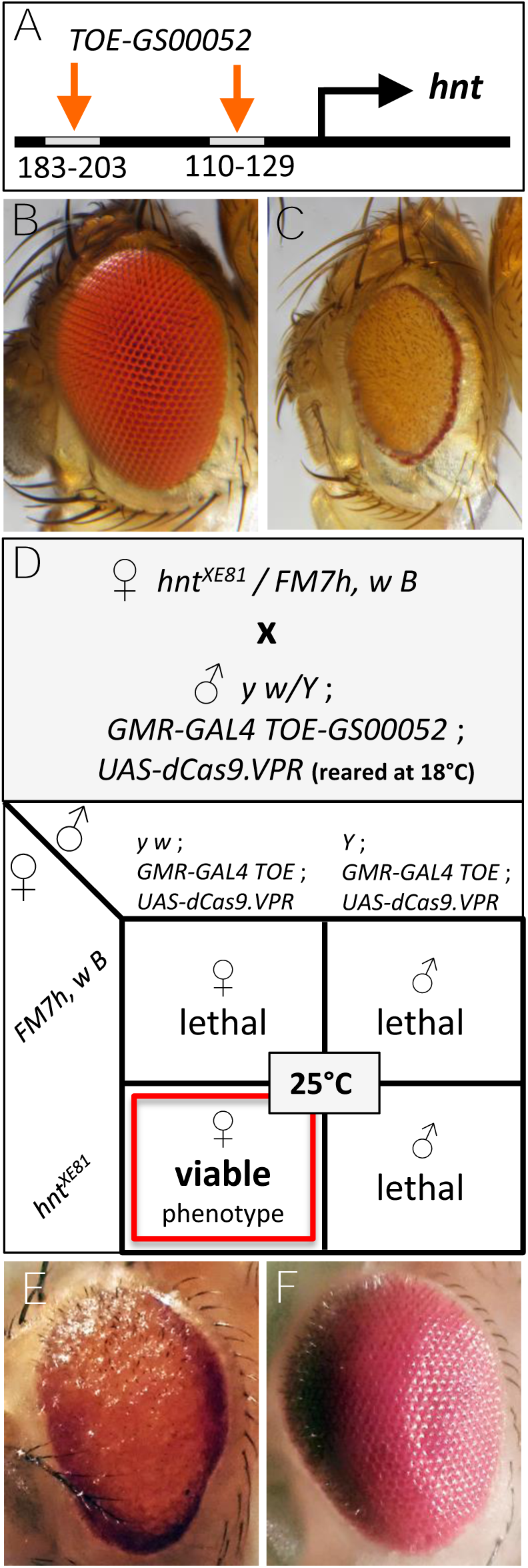
Overexpression of *hnt* disrupts eye development and the CRISPR/Cas9 mediated *hnt* overexpression phenotype is sensitive to temperature and gene dosage. (**A**) Schematic showing the sgRNA target sequences of *TOE-GS00052* relative to the TSS of *hnt*. (**B**) Wild type eye. (**C**) The *hnt* overexpression phenotype of *GMR-GAL4* > *UAS- GFP-Hnt*^*J27*^. **(D**) Nomenclature and associated Punnett square diagram of a balanced *hnt* allele (females) crossed to the CRISPR/Cas9 *hnt* overexpression stock (males). (**E**) Overexpression phenotype of *TOE-GS00052* + *GMR-GAL4*>*UAS-Cas9*.*VPR*. (**F**) Suppression of the overexpression phenotype *of TOE-GS00052* + *GMR-GAL4*>*UAS- Cas9*.*VPR* by co-expression of *UAS-hnt-RNAi*.

### Data Availability Statement

All Drosophila stocks used or recovered in this study are available upon request. The authors affirm that all data necessary for confirming the conclusions of the article are present within the article, figures, and tables.

## Results

Overall, we present our results as proof-of-principle for the concept that loss-of-function alleles in a GOI can be selected in an F1 screen. By way of example, in the following sections we describe our observations regarding *hnt* overexpression phenotypes, dosage sensitivity, the recovery of refractory *hnt* alleles, and the design and results of our novel screen for new *hnt* loss-of-function alleles.

### Characterization of dCas9.VPR Mediated Activation of *hnt* Expression

Previous work demonstrates CRISPR/Cas9 mediated *in vivo* activation of *hnt* expression. In this proof-of-principle example, ectopic *hnt* expression is shown in the third larval instar wing imaginal disc (Ewen-Campen *et al*. 2017). To further investigate CRISPR/Cas9 mediated overexpression of *hnt*, we used the *TRiP-TOE* insertion line *TOE-GS00052*, which ubiquitously expresses two sgRNAs, targeted to 110-129 bps and 183-203 bps upstream of the TSS of *hnt* (Fig. 1A). Crossing line *TOE-GS00052* to a line that carries the eye specific *GMR-GAL4* driver and *UAS-dCas9*.*VPR* resulted in pupal lethality at 25°C, but viable and fertile adults with a strong rough eye phenotype at 18°C. The *hnt* overexpression phenotype, which we have also observed in the context of *GMR- GAL4* > *UAS-GFP-Hnt*, is distinctive in that the eye margin is consistently more pigmented than the inner region, giving the appearance of dark rings around the eyes (Fig. 1 B,C). We confirmed that the rough eye phenotype in the context of *GAL4*>*UAS- dCas9*.*VPR* + *TOE-GSGS00052* is attributable to the overexpression of *hnt* by co-expression of *UAS-hnt-RNAi*, which resulted in a full suppression of the rough eye phenotype (Fig. 1E,F).

Through a series of crosses carried out at 18°C, we recovered a stock carrying all three transgenes required for CRISPR/Cas9 mediated activation of *hnt* expression in the pupal eye (i.e. *GMR-GAL4, TOE-GS00052*, and *UAS-dCas9*.*VPR*). When transferred to 25°C, this stock was 100% pupal lethal, with no adult escapers. To test the dosage responsiveness of *hnt* overexpression, we crossed males of the overexpression stock (reared at 18°C) to females carrying a balanced deletion of *hnt* (*hnt* is an *X*-linked gene, so the stock was *Df(1)ED6727/FM7*). Interestingly, at 25°C all progeny of this cross are *FM7*^*+*^ females and display the distinctive *hnt* overexpression rough eye phenotype, identical to *hnt*^*+*^/*hnt*^*+*^ overexpression phenotype at 18°C. We subsequently found that other *hnt* alleles previously characterized as strong loss-of-function alleles (*hnt*^*XE81*^, *hnt*^*1142*^, *hnt*^*345x*^) also behave the same way, and only produce *FM7*^*+*^ female progeny at 25°C (Fig. 1D). Other *hnt* alleles tested, including the hypomorphic semi-lethal allele *hnt*^*308*^, the temperature sensitive hypomorphic allele *hnt*^*peb*^, and the lethal allele *hnt*^*PL67*^, do not produce progeny at 25°C.

### The recovery of *hnt* alleles refractory to CRISPR/Cas9 mediated overexpression

The above results suggest that suppression of dCas9.VPR mediated activation of gene expression could be used not only to characterize existing alleles, but to select newly induced loss-of-function mutations in a GOI. To be broadly applicable, however, it is necessary to recover GOI alleles that are refractory to CRISPR/Cas9 mediated activation of gene expression, but are in all other respects wild type. In general, such “*GOI*^*REF*^” alleles permit the recovery of heterozygotes in situations where an overexpression phenotype is not sensitive to dosage or temperature. For example, in a CRISPR/Cas9 overexpression background, a *GOI*^*REF*^/*GOI*^*+*^ heterozygote might be associated with a visible phenotype or be lethal, whereas a *GOI*^*REF*^/*GOI*^*null*^ heterozygote would be incapable of GOI overexpression and would, therefore, have normal viability and no overexpression phenotype.

In order to recover *hnt*^*REF*^ alleles (*hnt* alleles refractory to CRISPR/Cas9 mediated activation of gene expression – but otherwise wild type), we used the approach of transmitting the ubiquitously expressed TOE-sgRNA transgene *GS00052* through the female germline with active Cas9 expression (crossing scheme is outlined in Fig. 2). In our case, we were able to cross such females to males carrying all three transgenes required for CRISPR/Cas9 mediated activation of gene expression of *hnt* in the pupal eye (*i*.*e*. the *y w*; *GMR-GAL4 TOE-GS00052*; *UAS-dCas9*.*VPR* stock that is viable at 18°C). Progeny of this cross that carried all three transgenes, but not displaying the overexpression phenotype were found in abundance. In order to recover isogenic refractory lines, single refractory males were crossed with *FM7* females, and through selection and backcrossing, isogenic refractory lines devoid of the autosomal transgenes were established (Fig. 2, crossing scheme 1). Two such *y w* lines were isolated and sequenced and both proved to carry deletions in both sgRNA target sites as compared to the control parental *y w* sequence (since *hnt* and *w* are tightly linked, we selected *y w hnt*^*REF*^ lines to ensure that we would be comparing the sequence of newly induced *hnt*^*REF*^ alleles with the sequence from the parental *y w* line). Other lines were established by backcrossing single refractory males to *y w*; *GMR-GAL4 TOE-GS00052 / CyO*; *UAS- dCas9*.*VPR / TM6* females (raised at 18°C). Subsequent crosses performed at 25°C allowed the selection of heterozygous viable *hnt*^*+*^/ *hnt*^*REF*^ females, which were again backcrossed, allowing for selection a triple homozygous stock: *hnt*^*REF*^; *GMR-GAL4 TOE- GS00052; UAS-dCas9*.*VPR*, from here on referred to as the “*RGV*” stock (Fig. 2, crossing scheme 2).

**Figure 2.**
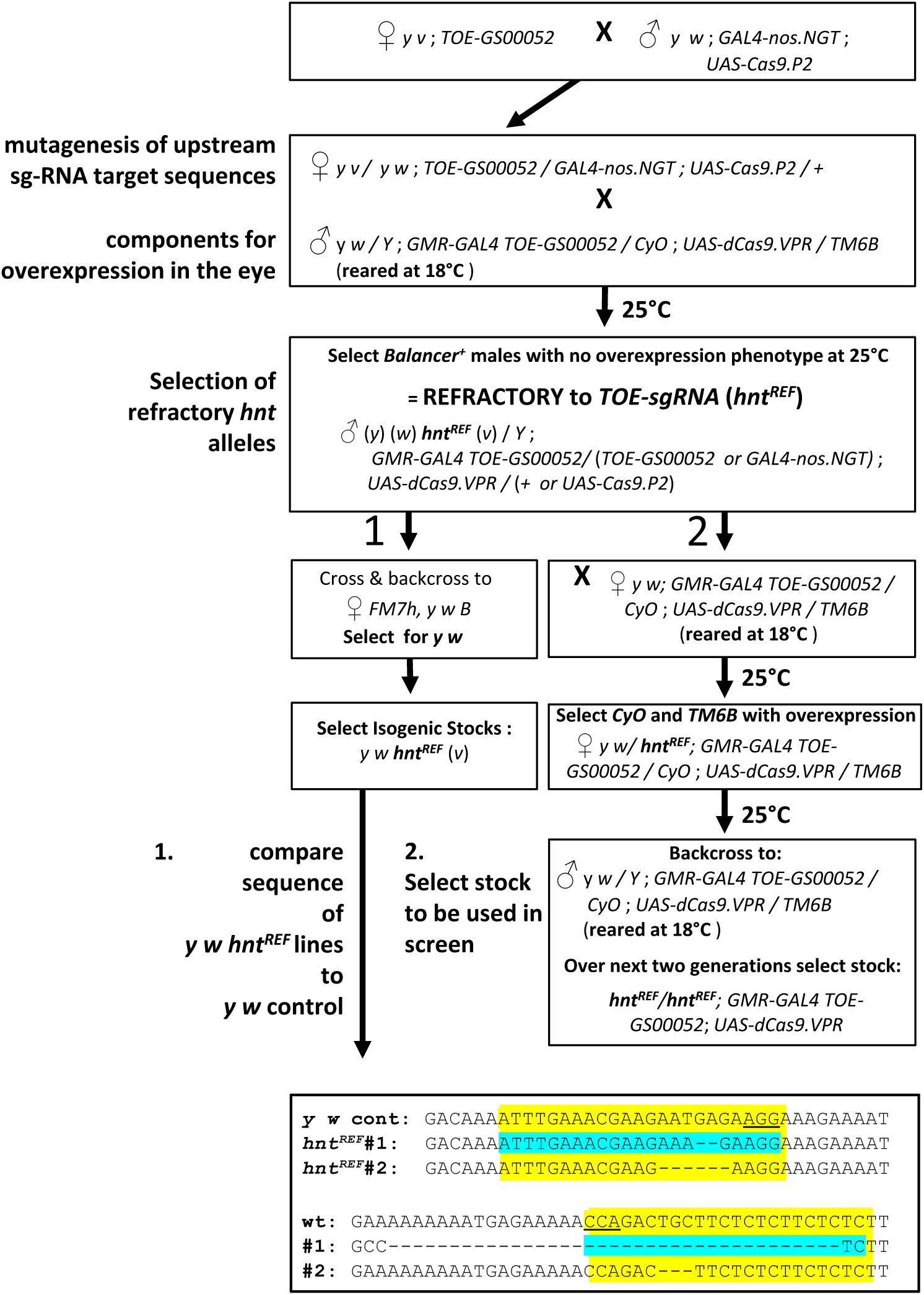
Creating chromosomes refractory to CRISPR/Cas9 mediated overexpression of *hnt*. Flow chart showing (**1**) the crossing scheme used to generate isogenic *hnt*^*REF*^ lines and the sgRNA target site sequences for two such lines; and (**2**) crossing scheme for the recovery of the *RGV* stock (*hnt*^*REF*^*/hnt*^*REF*^; *GMR-GAL4 TOE-GS00052*; *UAS- dCas9*.*VPR*). Selection of isogenic lines in crossing scheme (**1**) permitted the sequence of *hnt*^*REF*^ lines to be compared with the parental *y w* chromosome as *hnt* is tightly linked to *w*. Note that in crossing scheme (**2**) that the background markers on the *X* chromosome are not known and this stock is best described as (*y*) (*w*) *hnt*^*REF*^ (*v*). See Materials and Methods for full descriptions of stock genotypes.

We found the *RGV* stock to be viable with no overexpression phenotype raised at 25°C. We also checked the *RGV* stock by crossing females to *hnt*^*+*^ males (responsive to dCas9.VPR mediated activation of gene expression) and confirmed the expected phenotypes of viable female *hnt*^*+*^/*hnt*^*REF*^ progeny displaying the overexpression eye phenotype and viable male *hnt*^*REF*^ progeny with no overexpression phenotype. Likewise, we confirmed that *hnt*^*+*^/*hnt*^*+*^ females crossed to *RGV* males results in only female progeny that display the overexpression phenotype. In addition, we also retested *Df(1)ED6727* as well as the strong *hnt* alleles (*hnt*^*XE81*^, *hnt*^*1142*^, *hnt*^*345x*^) by crossing females of these *FM7* balanced lines to *RGV* males, which confirmed that *FM7*^*+*^ female progeny (*hnt*^*-*^/*hnt*^*REF*^) are viable and show no overexpression phenotype. Similar crosses to other *hnt* alleles (*hnt*^*peb*^, *hnt*^*308*^, and *hnt*^*PL67*^) confirmed that female heterozygous progeny (*hnt*^*-*^/*hnt*^*REF*^) are, in these cases, viable but do show the *hnt* overexpression phenotype.

### An F1 visible screen for recessive lethal mutations of *hnt* using the RGV stock

As a proof-of-principle, we used the *RGV* stock to screen for new mutant alleles of *hnt* induced by EMS mutagenesis. In the screen (outlined in Fig. 3), *RGV* virgin females were crossed to mutagenized males carrying the [*w*^*+*^] insertion *PBac{RB}e02388b*, which is tightly linked to *hnt* (inserted 6104 bp upstream) and has an unusual and easily scored crescent pattern of [*w*^*+*^] expression in the posterior eye. Recovery of newly induced *hnt* mutations was performed as an F1 visible screen for female progeny lacking the *hnt* overexpression phenotype. We screened approximately 45,000 progeny, from which we isolated 39 F1 females lacking the *hnt* eye overexpression phenotype. Of the 39 rare female progeny, 15 lines bred true and were established as balanced stocks over *FM7h,w* lacking the autosomal [*w*^*+*^] transgenes *GMR-GAL4* and *UAS-dCas9*.*VPR* from the *RGV* stock.

**Figure 3.**
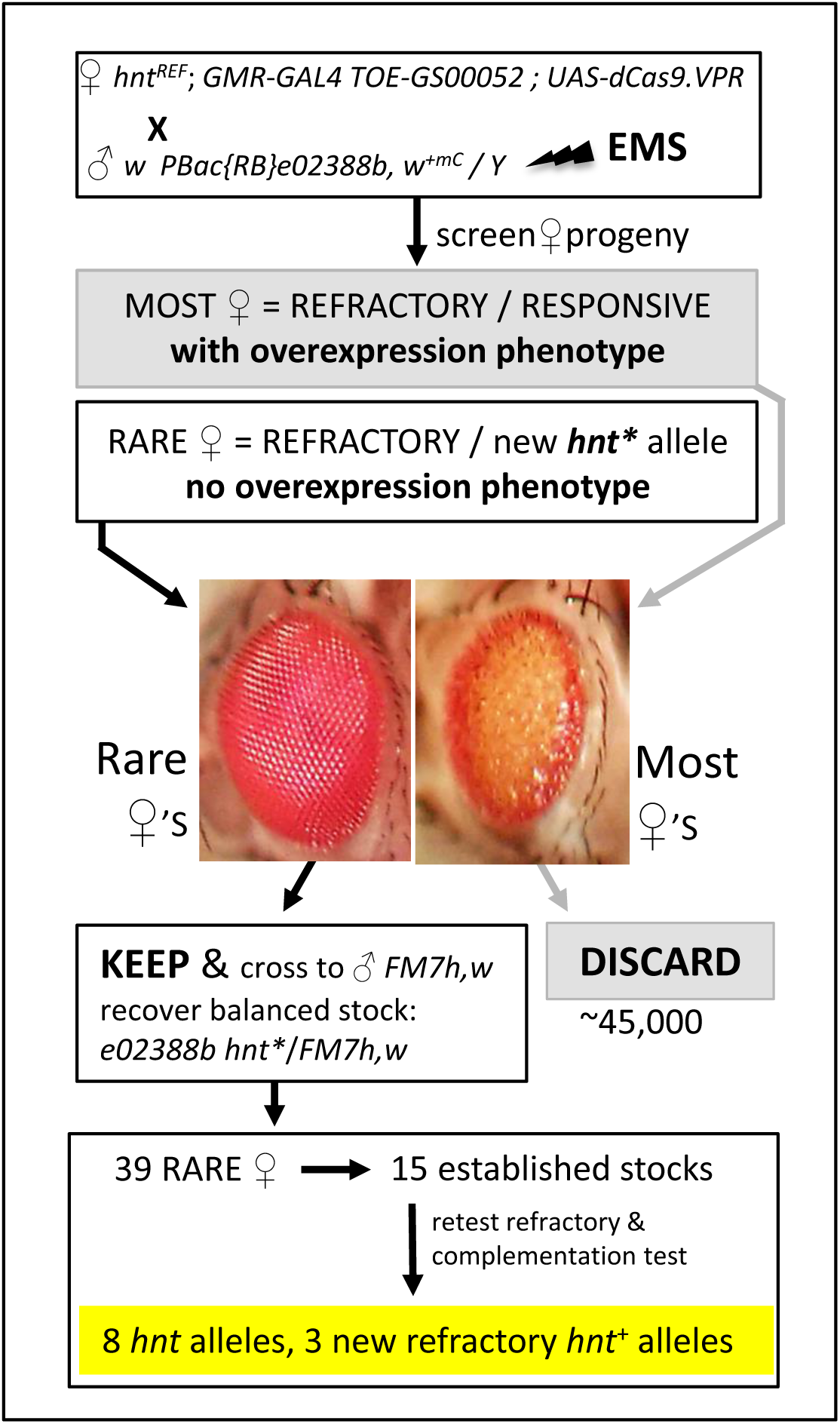
Crossing scheme and screen for the selection of EMS induced recessive lethal mutations of *hnt* as an F1 visible screen. Virgin females of the *RGV* stock crossed to EMS treated males results in the vast majority of females showing the *hnt* overexpression phenotype. Rare females not showing the *hnt* overexpression phenotype are selected as balanced stocks (*w PBac{RB}e02388b hnt*\* / *FM7h,w*) and tested to determine if they are new *hnt* loss-of-function alleles or newly induced refractory chromosomes.

### Characterization of new *hnt* alleles

Balanced lines recovered in the screen were retested for their refractory nature by crossing to *RGV* males and also tested for complementation with *hnt*^*XE81*^ (see Materials and Methods for description of complementation cross). Following retesting, we recovered eight new mutant alleles of *hnt* (all lethal alleles) and seven lines that carried new *hnt*^*REF*^ (*hnt*^*+*^) alleles. Of note, *hnt*^*REF-BHR7*^ was found to be mildly responsive to overexpression in males in the *RGV* retest, but completely refractory in females. All new *hnt*^*REF*^ alleles were sequenced by direct PCR sequencing (Fig. 4). The line *hnt*^*REF-BHR7*^ was found to have a single base pair change A >T within the first sgRNA target sequence, and *hnt*^*REF-WN72*^ was found to have a single base pair change of C >T within the PAM motif of the second sgRNA target sequence. Interestingly, these two *hnt*^*REF*^ alleles, where each is associated with a single base pair change affecting one sgRNA target sequence or the other, suggests that dCas9.VPR activation of *hnt* expression is only effective when both guide target sequences are fully intact and functional. All other *hnt*^*REF*^ lines were found to contain an identical deletion spanning from the first guide target to the second. Since each of these five lines was isolated from the same bottle, this likely corresponds to a “premeiotic cluster”, resulting from a mutation induced in a germline stem cell. Overall, therefore, we conclude that our screen resulted in the recovery of eight new *hnt* mutant alleles and only three new *hnt*^*REF*^ alleles.

**Figure 4.**
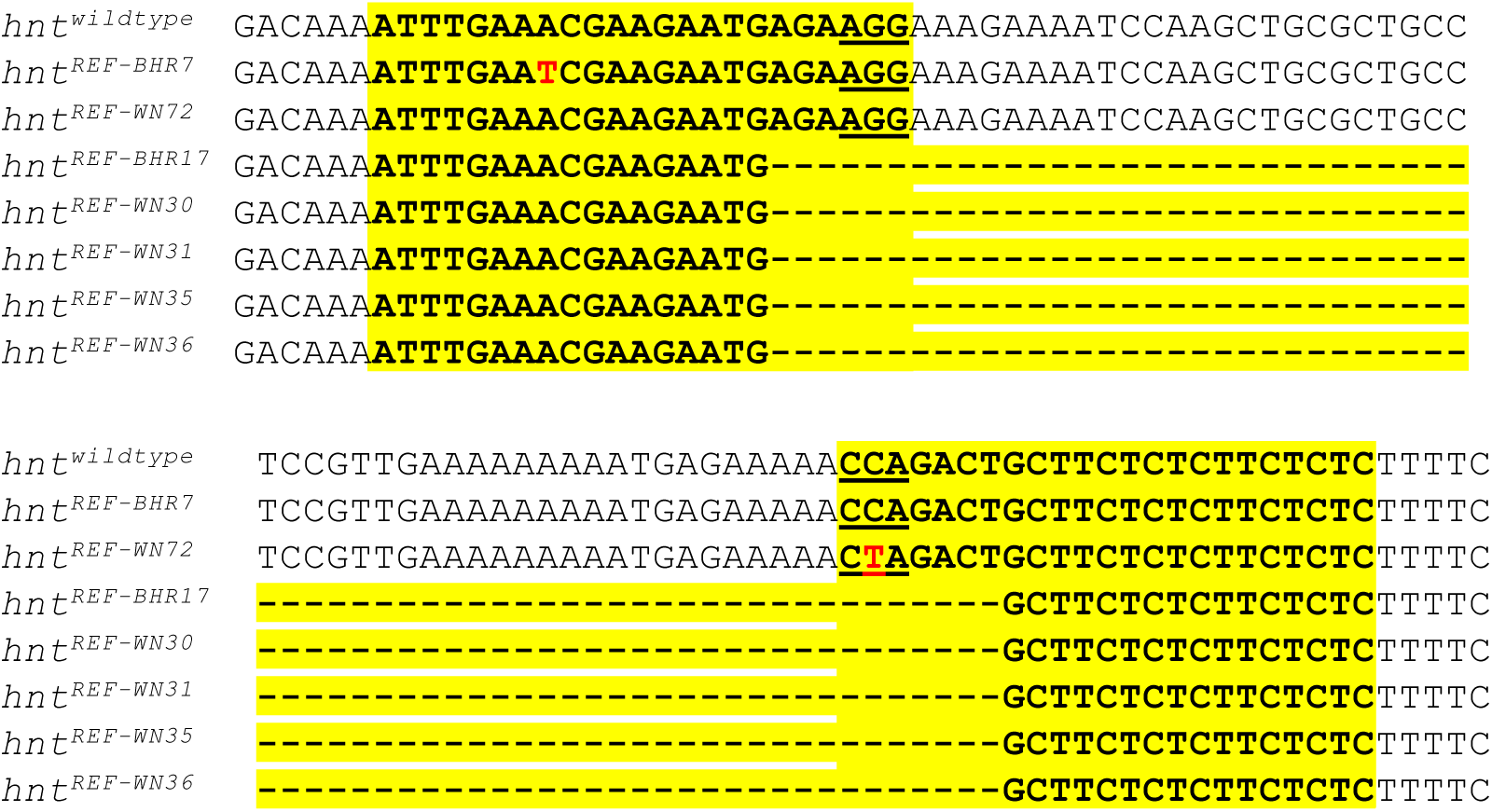
EMS-induced *hnt*^*REF*^ alleles are associated with mutations in the *sgRNA* target sequences of *TOE-GS00052*. Sequencing results for seven newly recovered EMS induced *hnt*^*REF*^ alleles shown in alignment with the reference genomic sequence (topmost). The *TOE-GS00052* sgRNA target sequences are highlighted in yellow. The PAM motifs for the two sgRNA targets sequences of *TOE-GS00052* are underlined. Single base pair changes of *hnt*^*REF-BHR7*^ and *hnt*^*REF-WN72*^ are shown in red font. The bottom five *hnt*^*REF*^ allele sequences are identical, indicative of a premeiotic mutation and represent a single mutagenic event.

All new *hnt* mutant alleles were also confirmed by failure to complement the temperature sensitive hypomorphic allele *hnt*^*peb*^ at the restrictive temperature of 29°C. Using *Ubi-DEcadherin-GFP*, we imaged pupal eyes of all *hnt*^*peb*^/*hnt*^*new*^ heteroallelic combinations at 25°C, which we found to be a more sensitive background for the *hnt*^*peb*^ phenotype. The severity of each allele was ranked by quantification of cones cells, which invariably number four per ommatidium in wild type (Fig. 5A). Mutant ommatidia frequently contain fewer than four cone cells per ommatidium (Fig. 5B-D, white arrowheads). Occasionally mutant ommatidia were found to contain 5 or 6 cone cells; these likely result from ommatidial fusion as they also contain three rather than two primary pigment cells (Fig. 5C, blue arrowheads). According to this analysis, none of the newly induced alleles is as severe as the *Df(1)ED6727* control, suggesting that none correspond to a null allele. We subsequently performed immunostaining on all lines using both monoclonal and polyclonal anti-Hnt antibodies. All alleles were found to be anti-Hnt positive using the polyclonal antibody (Figure 6, Table 1). We subsequently found that the strong hypomorphic EMS induced allele *hnt*^*XE81*^, previously described as antibody-null (using the same monoclonal antibody 27B8 1G9), also stains positive for Hnt using the polyclonal antibody (Fig. 6E). Interestingly, two alleles (*hnt*^*BHR49*^ and *hnt*^*WN52*^) were monoclonal negative but polyclonal positive, and although the polyclonal signal was expressed with correct tissue specificity, it did not show the normal nuclear localization of Hnt (Fig. 6D). These results further support the notion that none of these new *hnt* alleles corresponds to a null allele and that most EMS induced alleles of *hnt* produce non-functional protein.

**Table 1.**
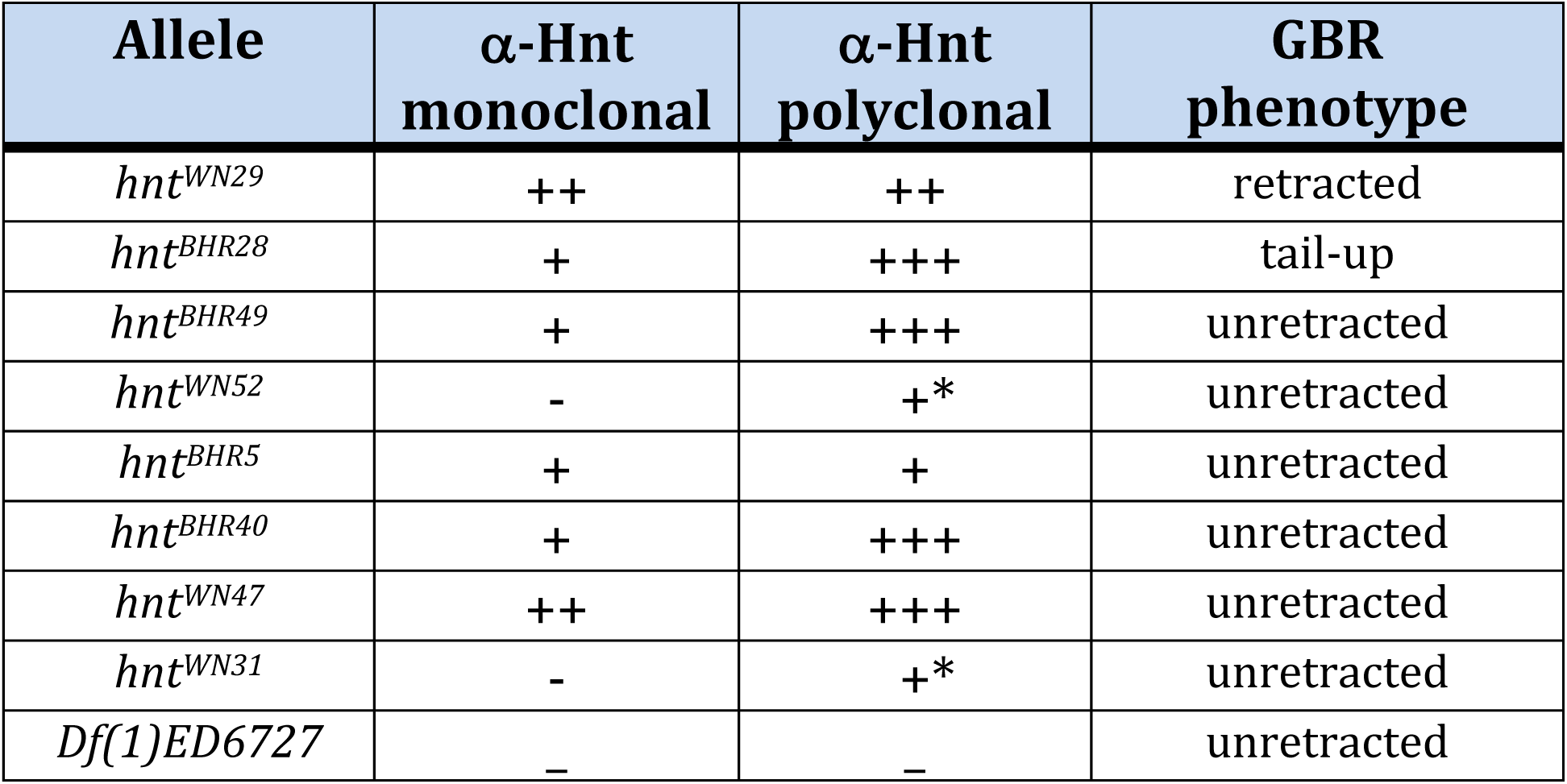
Summary of immunostaining results using anti-Hnt monoclonal and polyclonal antibodies as well as terminal phenotype analysis of new *hnt* alleles. * Anti-Hnt signal was not localized to the nuclei.

**Figure 5.**
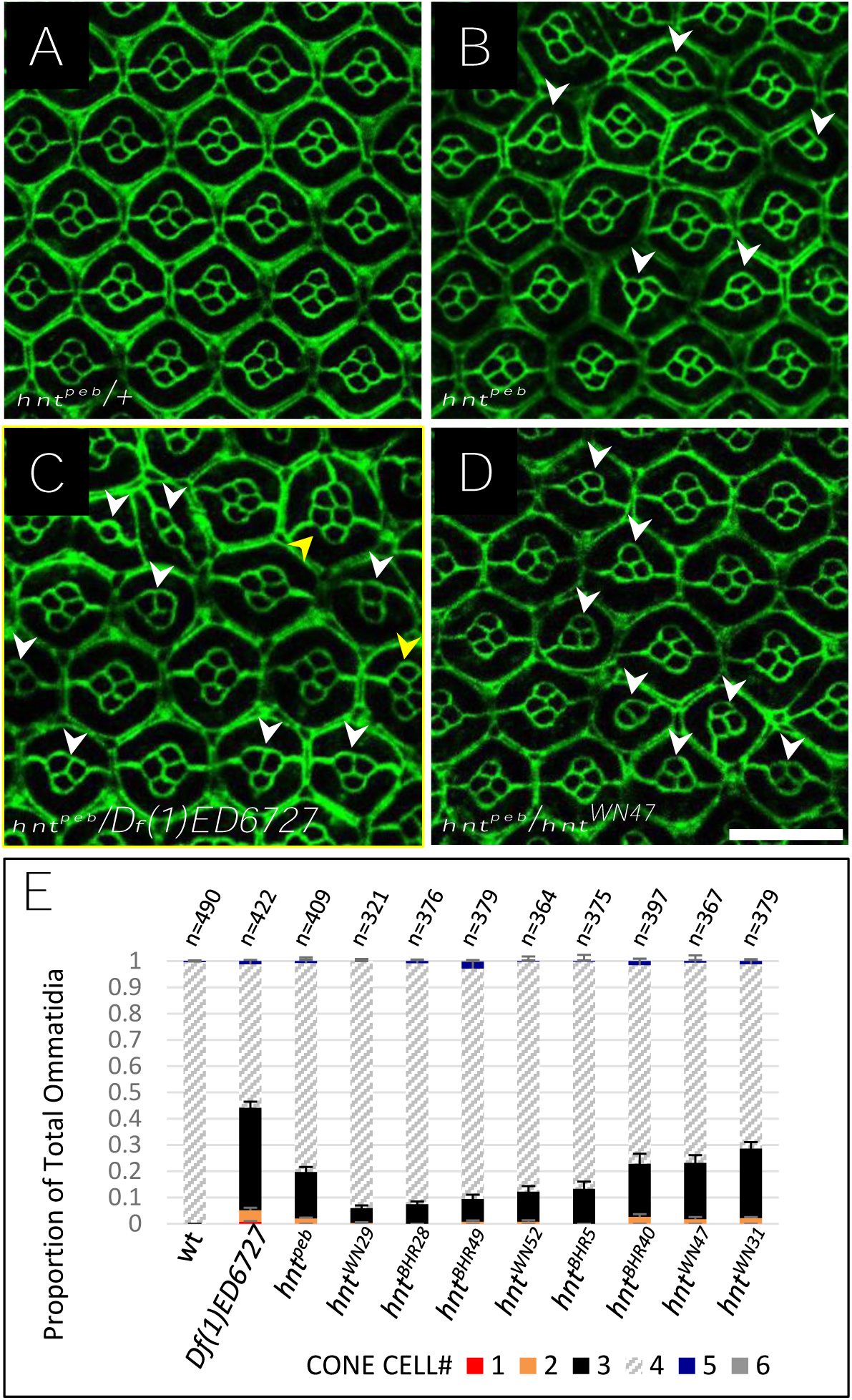
New *hnt* alleles can be arranged into an allelic series of strong hypomorphs according to their pupal eye phenotype when heterozygous to *hnt*^*peb*^. (**A**) *Ubi-DEcadherin-GFP* expression in control *hnt*^*peb*^*/+* pupal eye showing the normal ommatidial structure with four cone cells per ommatidium. **(B**) Homozygous *hnt*^*peb*^*/h*nt^peb^ reared at 25°C showing abnormal ommatidia having fewer than four cone cells (white arrowheads). (**C**) *hnt*^*peb*^/*Df(1)ED6727* raised at 25°C showing abnormal ommatidia with more (blue arrowheads) or less than four cone cells (white arrowheads). **(D**) *hnt*^*peb*^*/hnt*^*WN47*^ reared at 25°C showing abnormal ommatidia having fewer than four cone cells (white arrowheads). (**E**) Stacked bar graph showing the distribution of ommatidia cone cell numbers in wild type, and new *hnt* alleles heterozygous to *hnt*^*peb*^ reared at 25°C. Scale bar represents 20 μm.

**Figure 6.**
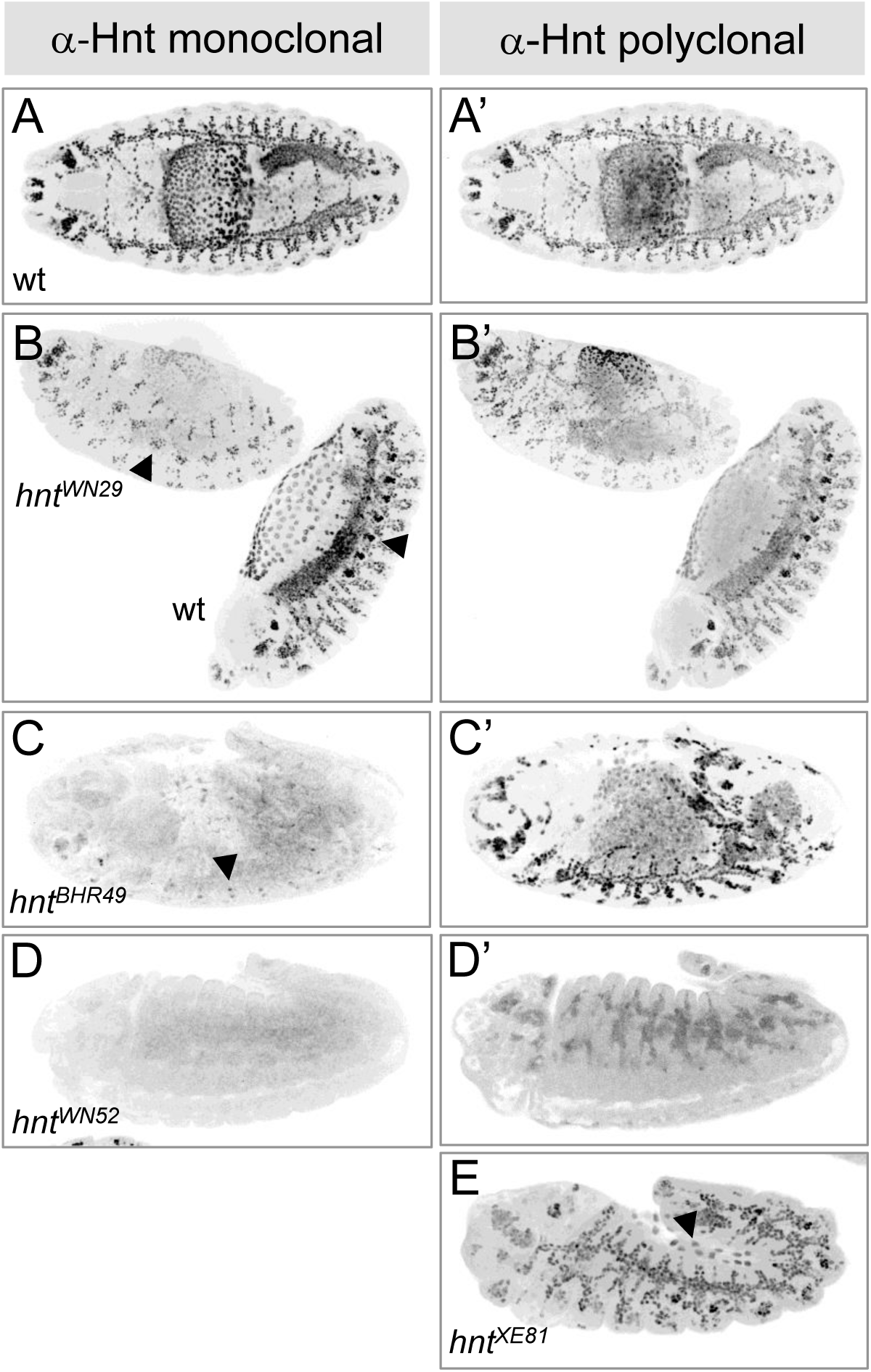
Examples of immunostaining of *hnt* mutants using anti-Hnt monoclonal and polyclonal antibodies. Shown are representative anti-Hnt monoclonal (**A-D**) and anti-Hnt polyclonal (**A’-D’, E**) confocal micrographs. (**A, A’**) Wild type (wt) showing normal pattern of anti-Hnt immunostaining in a stage 15 embryo. **(B**,**B’**) *hnt*^*WN29*^ shown in same frame as wt sib showing reduced anti-Hnt signal using both monoclonal and polyclonal antibody. The strong oenocyte signal observed in wild type is not seen in the *hnt*^*WN29*^ mutant (arrowheads). (**C**,**C’**) *hnt*^*BHR49*^ u-shaped mutant embryo showing weak monoclonal (arrowhead) but strong polyclonal signal. (**D**,**D’**) *hnt*^*WN52*^ u-shaped mutant embryo showing absence of signal with the monoclonal antibody and weak non-nuclear signal with the polyclonal antibody. (**E**) *hnt*^*XE81*^ u-shaped mutant with partially degenerated amnioserosa (arrowhead) showing strong anti-Hnt signal using the polyclonal antibody.

## Discussion

Overall, we have invented a new way to recover randomly induced mutations in a GOI based on CRISPR/Cas9 mediated activation of gene expression. As a proof-of-principle we applied this method to the gene *hnt* and recovered 8 new loss-of-function alleles. In addition to newly induced loss of function alleles, we also recovered newly induced refractory chromosomes in which the sgRNA target sequences had undergone mutation. While the recovery of new refractory alleles represents a source of false positives for this screening method, such false “hits” are easily recognized by complementation tests and sequencing the sgRNA target sites.

Considering the 8 new *hnt* alleles recovered, all showed a strong refractory phenotype in the cross to the *RGV* stock. However, we also isolated many F1 progeny that showed intermediate phenotypes. We were very interested in the possibility of recovering weak loss-of-function *<u>hnt</u>* alleles from these “hits”, but none of these lines bred true as X-linked mutations. We were also aware of the possibility of recovering dose-dependent second site modifiers. Two such lines were indeed recovered that consistently resulted in an intermediate dCas9.VPR mediated *hnt* overexpression phenotype, but do not map to the X chromosome. Further analysis of these lines may allow us to identify genes that are required for the full penetrance of the *hnt* overexpression phenotype. Such genes may ultimately shed light on the pathways and target genes regulated by Hnt.

### General Comments on Setting up an Overexpression Screen

As mentioned above, it is not an absolute requirement that a GOI show dosage sensitivity in order to design an overexpression screen, but it is useful to determine if this is the case. If the overexpression phenotype for a GOI is visible, with good viability and fertility, then one can determine if it is sensitive to gene dosage by crossing the overexpression background (*sgRNA* + *GAL4 driver* + *UAS-dCas9*.*VPR*) to a known loss-of-function allele or a deletion of the GOI and assaying for suppression of the visible phenotype. If such is the case, a screening strategy can be developed without using an allele that is refractory to dCas9.VPR mediated gene expression. If, however, the GOI is associated with a visible overexpression phenotype that is not sensitive to gene dosage, or if overexpression of a single responsive allele proves to be lethal, then an allele of the GOI that is refractory to dCas9.VPR mediated activation of gene expression is required. In this latter case, a refractory allele heterozygous to a loss-of-function or deletion of a GOI in the dCas9.VPR overexpression background will be viable and have no overexpression phenotype (lethal or visible).

### Can these Techniques be Generalized?

The generalization of our screening method will likely depend on the ability to recover refractory alleles of the GOI – that is, alleles whose only phenotype is that of not being responsive to CRISPR/Cas9 mediated activation of gene expression. Two questions remain to be answered before this technique can be generalized. First, is it always possible to create a refractory allele for a GOI, or is this only possible for certain genes? And second, if the overexpression phenotype for a GOI is not sensitive to temperature or gene dosage, would it still be possible to recover a refractory allele? The first question cannot be answered until CRISPR/Cas9 mediated activation of gene expression is used on more genes. In regards to the second question, it is clear that the development of our system was greatly enhanced by the striking temperature and gene dosage sensitivity of the *hnt* overexpression phenotype in the pupal eye. But what if just one copy of the GOI is sufficient to generate a lethal phenotype in the background of CRISPR/Cas9 mediated activation of gene expression (*sgRNA* + *GAL4* + *UAS- dCas9*.*VPR*)? Assuming the structure of a gene is such that it can be mutated to create a refractory chromosome, it should still be possible to select viable refractory alleles. This could be accomplished by a series of crosses in which a chromosome carrying a wild type allele in the GOI is put in the background of the TOE-sgRNA and germline expression of active Cas9. The potentially mutated chromosome can then be recovered as a stock without the active Cas9 but still carrying the TOE-sg-RNA transgene. Crossing such stocks to a background carrying a *Df* for the GOI along with the appropriate *GAL4* + *UAS-dCas9*.*VPR* to drive overexpression will allow selection of refractory alleles. Further studies will be required to determine if such techniques are feasible.

### Gene Overexpression as New Method for Characterizing Alleles

In general, the approach of dCas9.VPR mediated expression of mutant alleles can be considered as a new method for characterizing alleles. In a way, this is similar to Muller’s classical test for hypomorphic alleles through the synthesis of duplications containing a third copy of the mutant allele being tested. In this case, a third copy of a hypomorphic allele is expected to ameliorate the mutant phenotype compared to the phenotype of the homozygous euploid mutant (Muller 1932). In the case of dCas9.VPR mediated activation of gene expression, if an overexpression phenotype is suppressed in mutant heterozygote (*m*/*+*) relative to the wild type homozygote (*+*/*+*), then two possibilities arise: 1) the mutant allele may not encode a functional protein; or 2) the mutant allele may be refractory to overexpression through mutation or deletion of its sgRNA target sequences. Although these two possibilities may not be mutually exclusive, sequencing the sgRNA target sequences of a mutant allele could rule out the second possibility and thereby suggest that a particular allele is associated with a disruption of the protein-coding region of the GOI. The characterization of mutant alleles can be further improved if a refractory allele of the GOI is available *(+*^*REF*^) and the mutant is known to have intact sgRNA target sequences. In this situation, any phenotype resulting from CRISPR/Cas9 mediated activation of gene expression in the heterozygote (*+*^*REF*^/*m*), would suggest that the mutant allele is able to express a protein that is fully or partially functional. In this case, the mutant is more likely to be hypomorphic in nature, and could be associated with reduced gene expression or reduced protein function. Consistent with this, we found that the allele *hnt*^*308*^ was fully responsive to dCas9.VPR mediated activation of gene expression. This allele, which is associated with a *P* element insertion 226 base pairs upstream of the TSS of *hnt*, was previously characterized as hypomorphic and displays a lower level of *hnt* expression (Reed *et al*. 2001). The dCas9.VPR activation of *hnt*^*308*^ expression is consistent with the hypomorphic nature of this allele, but also indicates that the sgRNA target sequences on the *hnt*^*308*^ chromosome are intact and that the protein coding region can produce Hnt product that is sufficient to generate the overexpression phenotype. The same interpretations also apply to the allele *hnt*^*PL67*^, which is an enhancer trap line associated with embryonic lethality but is largely uncharacterized.

## Acknowledgements

Stocks obtained from the Bloomington Drosophila Resource Center (NIH P40OD018537) were used in this study. We are grateful to the BDRC as well as the Harvard Medical School for genetic stocks. We are grateful to H. Lipshitz (University of Toronto) for additional stocks and reagents. We thank C. Steele of Nikon Canada Inc. for assistance with microphotography. This work was supported by a grant to B.H.R. from the Natural Sciences and Engineering Research Council of Canada (NSERC RGPIN- 2015-04458).

